# The mechanics of a continuous self-assembling *s*urface-layer aids cell division in an archaeon

**DOI:** 10.1101/2025.02.04.636414

**Authors:** Sherman Foo, Ido Caspy, Alice Cezanne, Tanmay A.M. Bharat, Buzz Baum

## Abstract

The surface layer or “S-layer” is a planar lattice of glycosylated proteins that coats a wide range of archaea and bacteria instead of a classical cell wall or capsular polysaccharides, insulating them from the extracellular space and providing the cell membrane with physical support. Although the S-layer’s role as a mechanical support for the membrane might be expected to hinder cell division, we show that in rapidly dividing *Sulfolobus acidocaldarius* cells, the S-layer protein SlaA self-assembles into flexible lattice that helps flattens the cytokinetic furrow to accelerate ESCRT-III dependent cell division - a role that is especially important under conditions of mechanical stress. Taken together, these results generated using mutational analysis, live and fixed cellular imaging, along with electron cryomicroscopy, define the rules governing S-layer self-assembly and show how the mechanical properties of flexible lattice coats can enhance membrane functions to both physically support a cell and help to drive ESCRT-III dependent cell division.

**Significance statement:** The bounding membrane of a cell must be protected from environmental insults. In many bacteria and archaea that lack a cell wall, this is achieved by an enveloping S-layer. By virtue of its rigid, planar structure, the S-layer is also expected to act as a barrier to the cell shape changes required for cytokinesis. In this study of the S-layer lattice in the archaeon *Sulfolobus acidocaldarius* however, we demonstrate a role for this self-assembling mechanical support in re-shaping the cytokinetic furrow as an aid to ESCRT-III-mediated division. In showing how mechanically active and passive structures can work together to give rise to mechanically stable cells that can divide, this reconciles what seems like a trade-off between resilience and flexibility.

## Introduction

Because eukaryotes likely emerged from a merger between a member of the Asgard archaea and an alphaproteobacterial cell that gave rise to mitochondria (1, 2), many of the machines that drive core cell biological processes in eukaryotes have archaeal counterparts. For example, both human and *Sulfolobus acidocaldarius* cells use homologous machinery, consisting of ESCRT-III proteins and an AAA-type ATPase Vps4, to drive membrane remodelling during the final stages of cytokinesis (3, 4). At the same time, archaeal cell division is also likely influenced by unique biophysical aspects of archaeal cellular components, including peculiarities of the archaeal membrane and the highly glycosylated protein surface layer (S-layer) that fully envelopes most archaeal cells (5–8). Since these aspects of archaeal cell biology are not well understood, here, using *S. acidocaldarius* as a model system, we explore the influence of the Sulfolobus S-layer on ESCRT-III dependent cell division.

S-layers are two dimensional para-crystalline structures composed of S-layer protein monomers found on the surface of prokaryotes (5–9). With notable exceptions, including archaea that use pseudomurein, methanochondroitin or protein sheaths (7, 9, 10), and *Ignicoccus hospitalis* which lacks an S-layer entirely (11, 12), most archaea are enveloped by an S-layer. As the barrier separating an archaeal cell and its external environment, this S-layer serves a plethora of functions in different species: it provides mechanical support for the membrane (13, 14), physically and chemically isolates the cell from the external environment (15, 16), protects cells from predation (17), and facilitates the capture of low concentration nutrients from the environment (18, 19).

*S. acidocaldarius* is a member of the TACK (Thaumarchaeota, Aigarchaeota, Crenarchaeota, and Korarchaeota) archaea and is one of the most experimentally tractable archaeal relatives of eukaryotes - making it an ideal model system in which to study different aspects of archaeal cell biology. The structure of the *S. acidocaldarius* S-layer was recently solved (20, 21) to reveal a pseudocrystalline array of SlaA protein enveloping the cell surface with a p3 symmetry, bound to SlaB which functions as a membrane anchor (20–24). Both SlaA and SlaB proteins are highly glycosylated (21, 25, 26), a characteristic that has been suggested to play a role in guiding SlaA lattice assembly (21).

In bacteria and in halophilic archaea distantly related to *S. acidocaldarius*, live imaging experiments have revealed local insertion of S-layer monomers at the mid-cell during exponential growth, implying a mechanistic link between S-layer biogenesis and cell division in these organisms (27–30). In these systems, division occurs over an extended period of time, and is driven by the constriction of an FtsZ-based division ring in conjunction with new S-layer synthesis at the site of the furrow- analogous to the way cell wall synthesis aids cell division in some bacteria and yeast (31–33). This is very different however, to the situation in *S. acidocaldarius*, which divides rapidly using forces generated by changes in the state and preferred curvature of composite ESCRT-III polymers to deform and cut its bounding membrane (3, 34).

To investigate the assembly of the S-layer and its role in *S. acidocaldarius* cell division, we have combined molecular genetics, S-layer reconstitution, electron cryomicroscopy (cryo-EM), electron cryotomography (cryo-ET) and high temperature live cell imaging methods (34). Through these analyses we describe the incorporation of new SlaA protein into gaps in pre-existing S-layer lattices as they grow, and demonstrate the ability of S-layer lattices to self-assemble on the surface of live archaea cells through the extension of the SlaA lattice margins, via attachment to both SlaB and a novel thermopsin-like membrane protein. Interestingly, an analysis of the localization of both exogenously added and endogenous partial SlaA lattices revealed that the S-layer has a preference for the division bridge of constricting cells. Furthermore, a comparison of live cell imaging of control and *sla* mutant cells revealed that the S-layer is rigid enough to flatten the underlying membrane on either side of the furrow to accelerate cytokinesis, aiding division, especially in cells subjected to physical perturbation. Taken together, these data establish the nature of S-layer assembly in *S. acidocaldarius* and define a mechanical role for the S-layer in ESCRT-III dependent cell division.

## Results

### The S-layer is not essential in *Sulfolobus acidocaldarius*

To investigate the role of the S-layer in *S. acidocaldarius,* we generated mutant strains lacking either *saci2355* (Δ*slaA*), *saci2354* (Δ*slaB)*, or both components of the S-layer (Δ*slaAB*) using established genetic methods (35). In each case, the identity of the mutants was validated via genotyping the loci involved and by whole genome sequencing (Fig. S1A).

We first used fluorescently conjugated concanavalin A (ConA), a lectin which binds to glycosylated proteins (21, 25), to visualise the surface of these cells relative to the control strain (Fig. 1A). While levels of ConA fluorescence were substantially reduced in cells lacking either SlaA or SlaB relative to the control (an uracil auxotrophic background strain MW001), and were further compromised in the Δ*slaAB* double mutant (Fig. 1A and 1B), ConA was still able to label Δ*slaAB* cells. This shows that, while S-layer proteins make a significant contribution to the total surface glycosylation as expected (21, 25), there are likely a host of additional surface proteins or lipids which are subject to glycosylation that are still accessible to ConA once the S-layer has been removed. We also confirmed that SlaA and SlaB are major targets of the glycosylation machinery using mass-spectrometry analysis of a ConA pull-down from the wild-type strain DSM639 (Fig. S1B); and by Western blotting ConA pull-downs from extracts of cells expressing HA-tagged variants of SlaA and SlaB (Fig. S1C).

**Figure 1.**
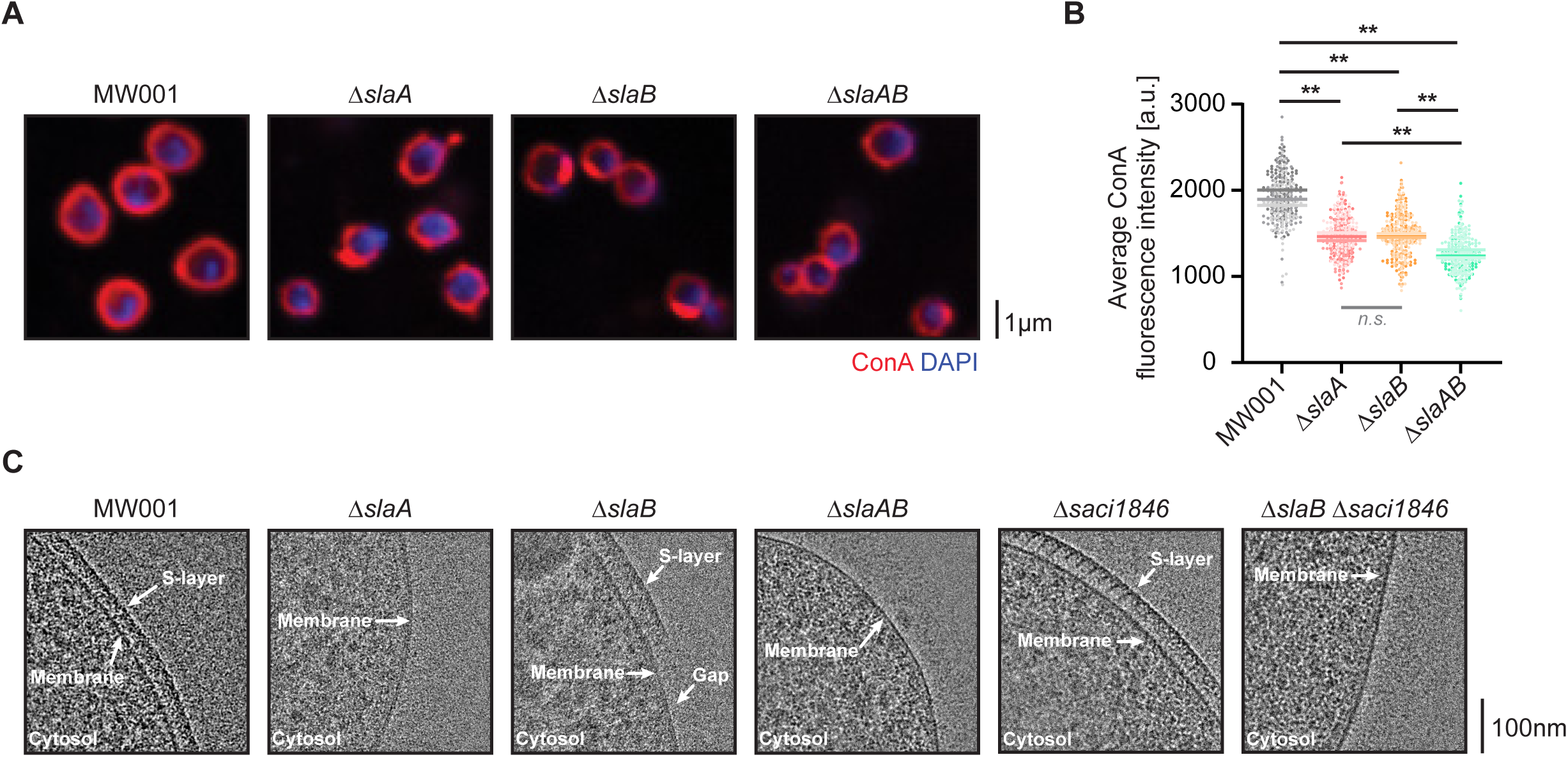
The S-layer of *S. acidocaldarius* is not essential for cell viability. (**A**) Markerless in-frame deletions of *slaA* (*saci2355*) and/or *slaB* (*saci2354*) genes were generated in the uracil auxotrophic strain MW001. Representative single-plane spinning-disk confocal images of these strains stained with the glycosylation marker ConA to visualise the highly glycosylated S-layer and DAPI to visualise DNA are shown. (**B**) Scatter plots of the quantification of the mutant strains show a decrease in average ConA fluorescence intensity in the deletion mutants (N=3, n=100 cells each). Each biological replicate is represented by a different shade and the means are indicated (MW001: 1911.4 ±71.4; Δ*slaA*: 1465.6 ±39.5; Δ*slaB*: 1478.7 ±18.8; Δ*slaAB*: 1260.5 ±35.9). *P*-values were derived using Welch’s *t*-test (N=3; \*\**p* ≤0.01; *n.s.*, no significance). (**C**) Representative cryo-EM images shows MW001 cells with an intact S-layer, Δ*slaA* and Δ*slaAB* mutants with no S-layer, and Δ*slaB* mutant with partial S-layer. A novel S-layer associated thermopsin-like protease protein Saci1846 assists SlaB in anchoring SlaA to the cell surface. Deletion of both *slaB* and *saci1846* results in the complete loss of the partial S-layer phenotype observed in the Δ*slaB* single mutant.

### A novel protein assists SlaB in anchoring SlaA to the cell surface

To obtain a higher resolution view of the surface of SlaA and SlaB mutant *S. acidocaldarius* cells, we next imaged cells using cryo-EM (Fig. 1C). In accordance with the recently published stalk (SlaB) and canopy (SlaA) model of the Sulfolobus S-layer (20, 21), removal of SlaA resulted in the complete loss of the S-layer (Fig. 1C, second panel). As expected, double mutant Δ*slaAB* cells resembled those of the Δ*slaA* mutant (Fig. 1C, fourth panel). Interestingly however, deletion of the membrane anchor SlaB gave rise to a discontinuous surface coat in which patches of intact SlaA lattices remained (Fig. 1C, third panel). This phenotype is consistent with observations in the SlaB mutant in the related species *Saccharolobus islandicus* and *Saccharolobus solfataricus* (36, 37), which led the authors to conclude that the SlaA S-layer peels away in cells that lack support of the membrane anchor SlaB.

Furthermore, in this cryo-EM analysis we were surprised to see that patches of S-layer remained at a reproducible distance of 37.2 ±1.3nm (n=6 tomograms) from the plasma membrane in cells lacking the membrane anchor SlaB (Fig. 1C, third panel); suggesting the presence of a redundant system. In *S. islandicus,* an S-layer associated protein M164_1049 was previously proposed to assist SlaB in stabilising the outermost SlaA layer (36). A pBlast search in *S. acidocaldarius* identified four homologs, Saci1846, Saci1049, Saci1534 and Saci2170. Of these, Saci1846, a thermopsin family protein, proved the closest homologue, being 36.95% identical to M164_1049 over nearly the entire length of the protein (Fig. S1D). To test if the *S. acidocaldarius* homologue of M164_1049 performs a conserved function in stabilising the S-layer, we generated mutants lacking Saci1846 in an MW001 or in a Δ*slaB* mutant background. Cells lacking Saci1846 only exhibited minor defects in the smooth S-layer structure, as visualised using cryo-EM (Fig. 1C and S1E). Yet, the S-layer remained anchored at a distance of 36.6 ±1.8nm (n=7 tomograms) from the plasma membrane in the Δ*saci1846* mutants (Fig. 1C, fifth panel). More strikingly, in line with previous reports from *S. islandicus* (36), the Δ*slaB* Δ*saci1846* double deletion mutant resembled the SlaA mutant in the entirely lacking an S-layer (Fig. 1C, sixth panel). Furthermore, trichloroacetic acid precipitation of proteins present in the growth medium of Δ*slaB* and Δ*slaB* Δ*saci1846* cultures (Fig. S1F) revealed abundant SlaA. These data indicate that SlaA is anchored at a similar distance of ∼37nm to the membrane via both SlaB and Saci1846, and is lost from the surface of *S. acidocaldarius* cells in their absence.

### S-layer biogenesis in *S. acidocaldarius*

Next, to investigate the dynamics of S-layer biogenesis and assembly, we attempted to identify sites of new S-layer insertion. Following optimization and validation of S-layer labelling using Alexa Fluor conjugated N-hydroxysuccinimide (NHS) ester dyes (Fig. S2A-C), we performed a pulse-chase experiment to compare the patterns of old and new S-layer on the cell surface. For this experiment, MW001 cells were first pulse-saturated with Alexa Fluor 488 NHS ester dye, and allowed to grow for 2h, before chase labelling was performed with Alexa Fluor 555 NHS ester dye. The cell surface was then examined by fluorescent microscopy. Interestingly, while some label was incorporated at random positions in the surface layer, there was no obvious correlation in the localisation of the pulse and chase labelling (Fig. S2D), even in dividing cells (Fig. S2D, bottom row). Instead, new S-layer insertion appeared to occur in a distributed stochastic manner across the cell surface, likely filling in small gaps in the lattice that formed as cells grow. This is markedly different to the case reported in some organisms which grow via polar S-layer biogenesis such as *Caulobacter crescentus* (30), and is strikingly different from the case in organisms like Haloferax, where new S-layer deposition occurs at the site of division (29).

### Visualising the self-assembly of the SlaA S-layer

As an additional method by which to measure S-layer protein assembly, we designed a C-terminally tagged construct that could be used to express SlaA-HA under the control of an arabinose inducible promoter. Induction of the SlaA-HA expression in MW001 led to the seemingly random insertion of newly expressed SlaA-HA into the S-layer (Fig. 2A, left panel), in line with the pattern of insertion described above. Interestingly, however, when this experiment was repeated in the Δ*slaA* mutant, the newly expressed proteins formed islands on the cell surface that led to partial coverage of the cell surface (Fig. 2A, right panel). We hypothesized that this partial S-layer coverage phenotype observed in the Δ*slaA* SlaA-HA rescue strain might reflect the ability of S-layer proteins to spontaneously assemble into a lattice. This ability to self-assemble would enable newly exported SlaA-HA protein to diffuse along the cell surface, filling in gaps in the lattice that form during growth or as the result of mechanical disruption to yield the even and complete coverage of the membrane by the S-layer (Fig. 2B, left panel). In the Δ*slaA* SlaA-HA strain, by contrast, SlaA-HA was lost from the cell unless it was able to associate with an existing S-layer lattice. As a consequence, patches of SlaA-HA expanded outwards through self-assembly until they coated the entire cell surface (Fig. 2B, right panel).

**Figure 2.**
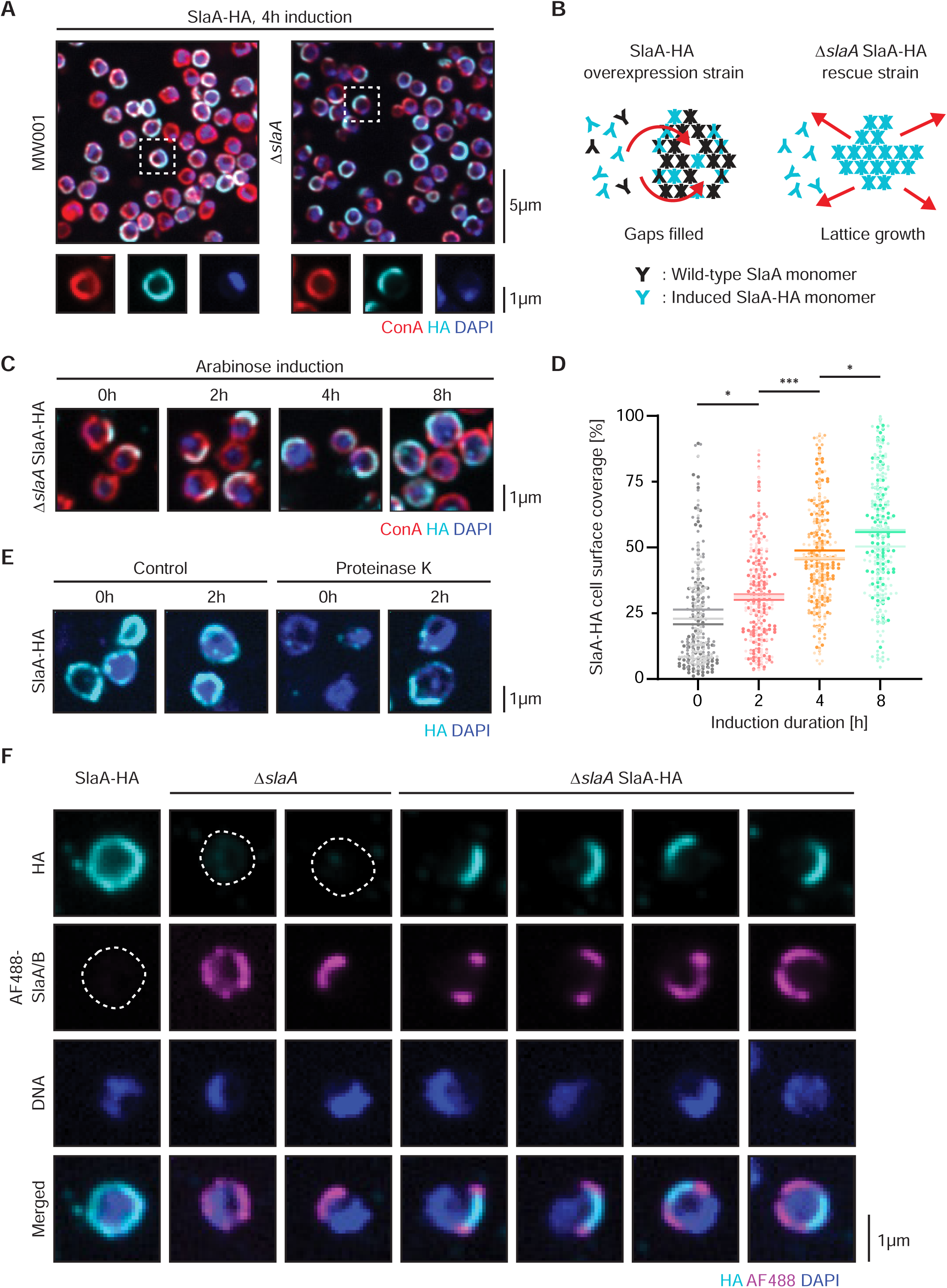
Visualising the SlaA S-layer biogenesis and assembly in *S. acidocaldarius*. (**A**) Field of view showing a representative single-plane spinning-disk confocal image of C-terminally HA-tagged SlaA following 4h of induction in the MW001 background strain (left) or Δ*slaA* deletion mutant (right). Magnified images of a representative cell indicated by the white box in each example are shown below. Overexpression of SlaA in MW001 displays evenly distributed HA-tagged S-layer protein on the cell surface. Rescue strain expressing HA-tagged SlaA in the Δ*slaA* deletion mutant background however displays a phenotype where the proteins are often localized to a crescent shaped patch on one side of the cell. (**B**) Model of the SlaA S-layer lattice formation in *S. acidocaldarius*. Expression of endogenous and HA-tagged SlaA in the overexpression strain ensures even coverage of the cell surface due to filling in of gaps in the lattice. In the rescue mutant, the SlaA-HA lattice self-assembles and expands outwards from a nucleation point. (**C**) Time course of representative single-plane spinning-disk confocal images of the rescue strain expressing HA-tagged SlaA following 8h of arabinose induction. (**D**) Quantification of the SlaA S-layer coverage as a percentage of the total cell surface at the middle z-plane shows an increase in cell surface coverage over time, supporting a nucleation model in the SlaA S-layer lattice assembly (N=3, n=100 cells each). Each biological replicate is represented by a different shade and the means are indicated (MW001: 23.3 ±2.4%; Δ*slaA*: 31.1 ±0.9%; Δ*slaB*: 46.8 ±1.5%; Δ*slaAB*: 54.3 ±2.8%). *P*-values were derived using Welch’s *t*-test (N=3; \**p* ≤0.05; \*\*\**p* ≤0.001). (**E**) Proteinase K treatment of HA-tagged SlaA overexpression strain shows a similar phenotype to the rescue mutants with partial S-layer coverage on one side of the cell upon removal and subsequent recovery of S-layer protein levels. Shown here are representative immunofluorescence images. (**F**) Addition of fluorescently labelled (Alexa Flour 488, AF488), exogeneous S-layer proteins to cell culture medium was performed to identify sites of S-layer growth. Representative immunofluorescence images shows that cells with an intact S-layer are not labelled with exogenous S-layer proteins, while cells lacking SlaA are. SlaA rescue mutants are labelled with exogenous S-layer proteins at the tips of the endogenous HA-tagged S-layer patches, demonstrating the self-assembly of new S-layer lattices in *S. acidocaldarius*.

To better probe this process of self-assembly, we next followed the induction of SlaA-HA in Δ*slaA* mutant cells over the course of 8 hours. An increase in the coverage of the cell surface with SlaA-HA was observed with time (Fig. 2C and 2D). To verify that this mode of assembly is not peculiar to the Δ*slaA* mutant, MW001 cells were treated with Proteinase K to remove the endogenous S-layer, prior to the induction of SlaA-HA expression (Fig. 2E). Again, in this case, the recovery of the S-layer was via nucleation and lattice growth, supporting this model of SlaA assembly (Fig. 2B).

These experiments using HA-tagged SlaA protein assayed the ability of newly expressed protein to be inserted into the S-layer following its export across the membrane via the translocon. To determine whether pre-folded SlaA protein can also be incorporated into a growing S-layer lattice, we attempted to coat cells with exogenous labelled SlaA. For this experiment, S-layer proteins were extracted from wild-type *S. acidocaldarius* cells using a slightly modified version of previously published protocols (Fig. S2E, see materials and methods) (21, 38). The purified S-layer proteins were then covalently conjugated with a fluorescent NHS ester dye and incubated with S-layer mutant strains. As assayed by flow cytometry, this method efficiently labelled the surface of Δ*slaA* mutant cells, where the membrane anchor SlaB is likely free to capture exogenous soluble SlaA, but not in Δ*slaB* mutants or in control cells that already possess a complete SlaA coat (Fig. S2F). When visualized using light microscopy, fluorescently conjugated exogenous SlaA was seen forming domains on the surface of Δ*slaA* mutant cells, but was not seen associating with the SlaA-HA overexpression strain, which likely lacks free surface SlaB (Fig. 2F). Strikingly, when exogenous labelled proteins was added to Δ*slaA* SlaA-HA strains expressing relatively low levels of HA-tagged SlaA, the labelled SlaA was recruited to the edges of the SlaA-HA patches as expected if the endogenously forming lattice is able to extend at its margins via the recruitment of pre-folded SlaA. Taken together, these data demonstrate that a complete S-layer in *S. acidocaldarius* is likely generated and maintained during cell growth via the energetically-favourable process of self-assembly in a manner which might not require local SlaA translation and export.

### The S-layer influences cell division in *S. acidocaldarius*

Previous studies in related Sulfolobales have implicated the S-layer in cell division (36, 37). To test whether or not SlaA plays an analogous role in *S. acidocaldarius*, we performed flow cytometry analysis to assess the DNA content of our panel of S-layer deletion mutants during exponential growth. At first glance, the profiles look similar. Populations of exponentially growing cells in all conditions have a relatively similar 1N and 2N DNA content, corresponding to the G1 and G2 phases of the cell cycle (Fig. 3A). In populations of cells lacking either SlaA and/or SlaB, however, we observed a significant, if slight, increase in the proportion of cells containing an abnormal >2N DNA content (Fig. 3A and 3B) – a phenotype indicative of a mild division defect.

**Figure 3.**
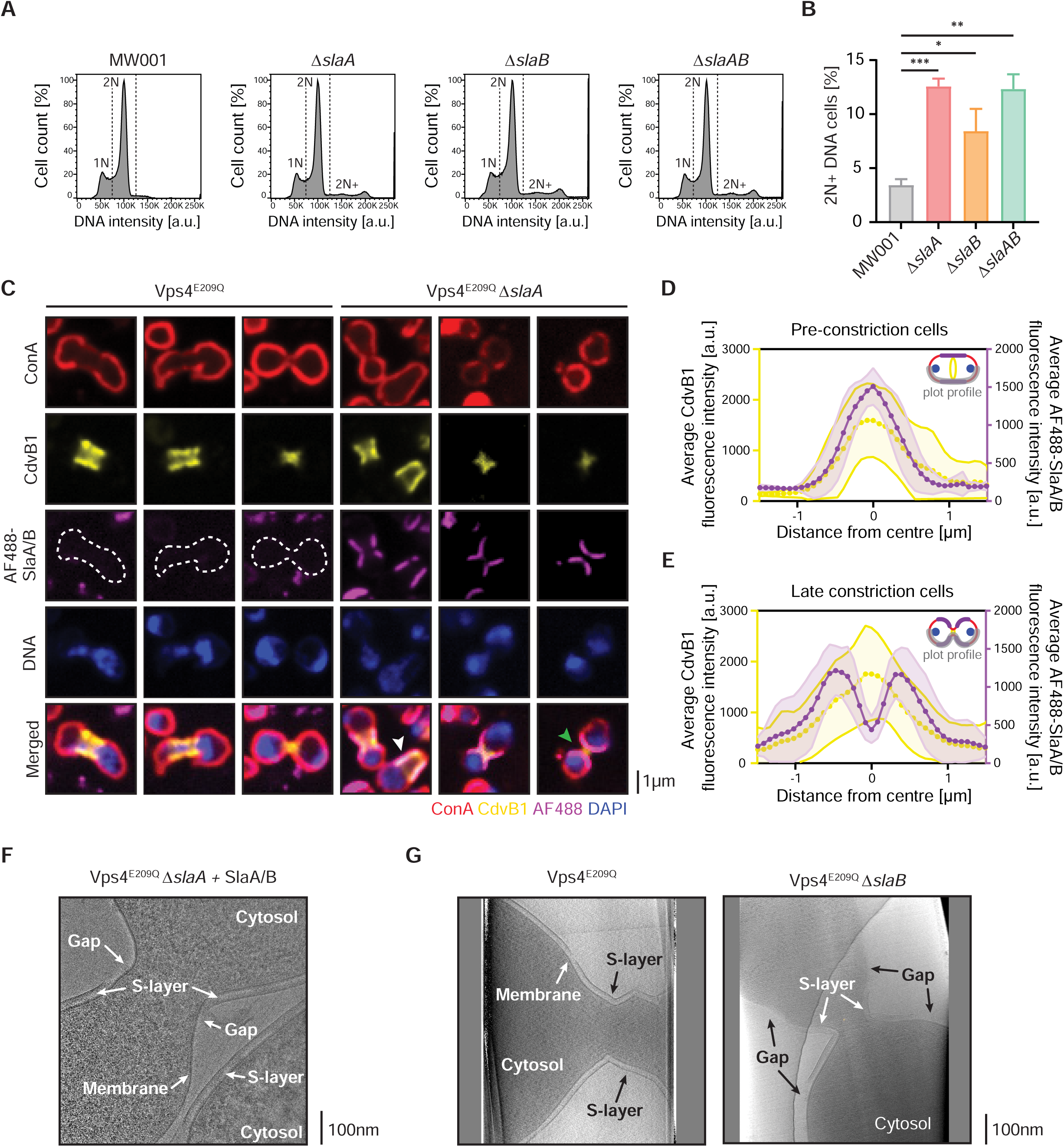
The S-layer is implicated in cell division and localizes preferentially at the division bridge. (**A**) Representative flow cytometry histograms of the DNA content in asynchronous cultures of MW001 and S-layer mutants. Most cells contain either 1N or 2N DNA content corresponding to the G1 or G2 phases of the cell cycle. Deletion of either SlaA and/or SlaB results in the accumulation of cells with abnormal 2N+ DNA content. (**B**) Quantification of percentage of cells exhibiting more than 2N DNA content in the flow cytometry experiments (N=3, n=5.0×10^5^ events). Shown here are means ±SD (MW001: 3.4 ±0.5%; Δ*slaA*: 12.6 ±0.6%; Δ*slaB*: 8.4 ±1.7%; Δ*slaAB*: 12.3 ±1.1%). *P*-values were derived using Welch’s *t*-test (N=3; \**p* ≤0.05; \*\**p* ≤0.01; \*\*\**p* ≤0.001). (**C**) Representative immunofluorescence images of MW001 and the Δ*slaA* mutant overexpressing the Walker B dominant negative mutant Vps4^E209Q^. Addition of fluorescently labelled, exogeneous S-layer proteins results in the binding of exogenous SlaA S-layer proteins to the division bridge of Vps4^E209Q^ Δ*slaA* mutants (95 ±0.82% of dividing cells with signal at the division bridge; N=3, n=100 cells) but not the Vps4^E209Q^ mutants. Shown here are representative single-plane spinning-disk confocal images. Plot profile of the average intensities of CdvB1 (yellow) and fluorescently labelled, exogeneous S-layer proteins (magenta) in pre-constriction cells (**D**) or late constriction cells (**E**). A diagram showing the plot profile used indicated in grey is shown. Shown here are means, and the SD are indicated by the shaded area (n=25 cells each). Examples of pre-constriction and late constriction cells are indicated by the white and green arrowheads respectively in (**C**). While fluorescently labelled, exogeneous S-layer proteins localizes to the middle of both pre and late constriction cells as marked by the ESCRT-III protein CdvB1, they are excluded from the middle where local curvature is highest in late constriction cells. (**F**) Representative cryo-EM image of Vps4^E209Q^ Δ*slaA* mutants incubated with unlabelled exogeneous S-layer proteins shows the formation of the S-layer lattice on the lateral sides of the division bridge. See also Fig. S3F. (**G**) Representative cryo-ET slices of MW001 and the Δ*slaB* mutant overexpressing the Walker B dominant negative mutant Vps4^E209Q^. MW001 expressing Vps4^E209Q^ has an intact S-layer encompassing the entire cell, while the Δ*slaB* mutants with a partial S-layer phenotype shows a preference for the SlaA S-layer localization at the division bridge (71.43% of cells with S-layer at the division bridge; n=21 cells). See also Fig. S3G.

These observations led us to explore the role of the S-layer in the shape changes that accompany cytokinesis more directly. To image division bridges in detail, we arrested control and *sla* mutant cells at different stages of division using a dominant negative Vps4 mutant with a E209Q point mutation in the Walker B motif (Vps4^E209Q^) that blocks ATP hydrolysis and ESCRT-III ring remodelling (3) (Fig. S3A). Exogenous labelled SlaA was then added to these cells to probe for gaps in the S-layer (Fig. S3B). In Vps4^E209Q^ control cells that express endogenous SlaA, the cells were not labelled by the addition of exogenous labelled S-layer proteins, implying the absence of gaps in the lattice that could be filled by exogenous protein, even in the division bridges where membrane remodelling occurred (Fig. 3C, left panels). By contrast, exogenously labelled S-layer proteins formed extended lattices on Vps4^E209Q^ Δ*slaA* cells, which preferentially coated division bridges (Fig. 3C, right panels) (95.0 ±0.82% of dividing cells with signal at the bridge; N=3, n=100 cells).

To test if this local accumulation of labelled protein might reflect the S-layer lattice’s preference for membranes with a defined curvature, we took advantage of the fact that Vps4^E209Q^ Δ*slaA* cells assume a variety of shapes to compare the labelling of cells in early constriction stage, where they take up an extended oval shape, to dumbbell shaped cells in late constriction stage. When we examined Vps4^E209Q^ Δ*slaA* cells arrested prior to furrow formation, the bias was striking. Plot profile of the cell surface shows that S-layer lattices were visible spanning the entire midzone of the cell, marked by the ESCRT-III homolog CdvB1 ring, but were absent from cell tips where curvature is high (Fig. 3D and S3C). This distribution was confirmed by a correlation analysis, which revealed a preference of the exogenous labelled S-layer for regions of lower curvature - something surprising given the overall curvature of *Sulfolobus* cells (Fig. S3D). Next, we looked at the localisation of exogenous SlaA in dumbbell shaped Vps4^E209Q^ Δ*slaA* cells. In this case, exogenous S-layer accumulated at the division bridge, but tended to be excluded from the very centre of the bridge, where the local curvature is highest (Fig. 3E and S3E). A cryo-EM analysis first confirmed that the exogenous-added labelled SlaA was able to form an S-layer on the surface of Δ*slaA* (Fig. 3F), which is very similar to that seen in the control (Fig. 1C). Moreover, this analysis showed that, while soluble S-layer proteins were able to form relatively flat lattices on the surface of Vps4^E209Q^ Δ*slaA* cells, the S-layer lattice tended to be absent from regions of the division bridge exhibiting very high local curvature, or appeared less uniform in these regions (Fig. 3F and S3F).

To test whether or not cells with an endogenous partial S-layer also preferentially localizes to division bridges, we next looked at unlabelled Δ*slaB* cells which retains patches of SlaA (Fig. 1C, third panel), expressing Vps4^E209Q^. Again, we observed a similar bias by cryo-ET (Fig. 3G and S3G, see also supplementary movies SM1-SM4). While a normal intact S-layer was observed enveloping the entirety of control Vps4^E209Q^ cells, in Vps4^E209Q^ Δ*slaB* cells, the patches of endogenous SlaA S-layer were preferentially localized at the division bridge (71.43% with S-layer at the division bridge; 15/21 cells). Unlike exogenous labelled S-layer however, the S-layer in Vps4^E209Q^ Δ*slaB* cells tended to remain continuous across the middle of constricted bridges and was only excluded from the central region in a relatively small fraction of cases (23.81%- 5/21 cells; 1/21 had no S-layer).

Taken together these data show that, while the S-layer remains intact at the division site of wild-type cells with few gaps, in cells with extended division bridges that have a partial S-layer, it preferentially coats and flattens the membrane at the division bridge and, when exogenously added, preferentially binds to regions of lower curvature.

### Cell division phenotypes of S-layer mutants

Having shown that the S-layer has a preference for sites of specific curvature and for division bridges, we wondered if the S-layer may affect the division machinery located at these division bridges. Intriguingly, the loss of the S-layer in Δ*slaA*, Δ*slaB* and Δ*slaAB* mutants did not visibly alter the localization of the division machinery itself, nor the stepwise changes in the composition of the ESCRT-III rings (CdvB, CdvB1 and CdvB2) driving division (4, 39–42), when assayed using either via flow cytometry (Fig. S4A) or immunofluorescence (Fig. S4B).

To determine if the S-layer plays a more direct mechanical role in cytokinesis instead, we imaged control MW001 cells and *sla* mutant cells labelled with CellMask Plasma Membrane stain as they grew and divided. Under these conditions, we noted that control MW001 cells had an irregular but near spherical shape during interphase, but a much more angular shape with a flat division furrow during cytokinesis (Fig. 4A). This was due to the fact that, as previously reported (34), the furrow in MW001 cells deepens at a relatively constant rate in a process that resembled slicing, to produce two daughter cells of similar sizes within a period of 10 minutes (Fig. 4A) (3, 43). The contrast with division in cells lacking SlaA or SlaB was striking: *sla* mutant cells were nearly spherical in interphase and remained round or dumbbell shaped throughout the process of cell division (Fig. 4B and 4C). This difference was significant as shown by the quantification of local curvature in regions of the cell close to the division bridge (Fig. 4D). Moreover, at intermediate timepoints, they exhibited smoothly curved division bridges. To assay the role of the S-layer in cytokinesis, we measured the time taken from the beginning until the end of cytokinesis. This revealed a dramatic and significant reduction in the constriction rate of the furrow in the absence of a complete S-layer (Fig. 4E). This was accompanied by an increase in the rates of division error - such as markedly asymmetric divisions (Fig. 3A). Finally, we subjected cells to osmotic shock as a way of physically perturbing them to determine whether the S-layer performs an important mechanical function under more physically challenging conditions. While division failures were rare in control cells shifted to hyperosmotic media containing 4% sucrose, a significant number of division bridges collapsed at late stages just prior to abscission in mutant cells lacking the S-layer (Fig. 4F and 4G) - as expected if abscission was facilitated by physical flattening of the cytokinetic furrow by the overlying S-layer.

**Figure 4.**
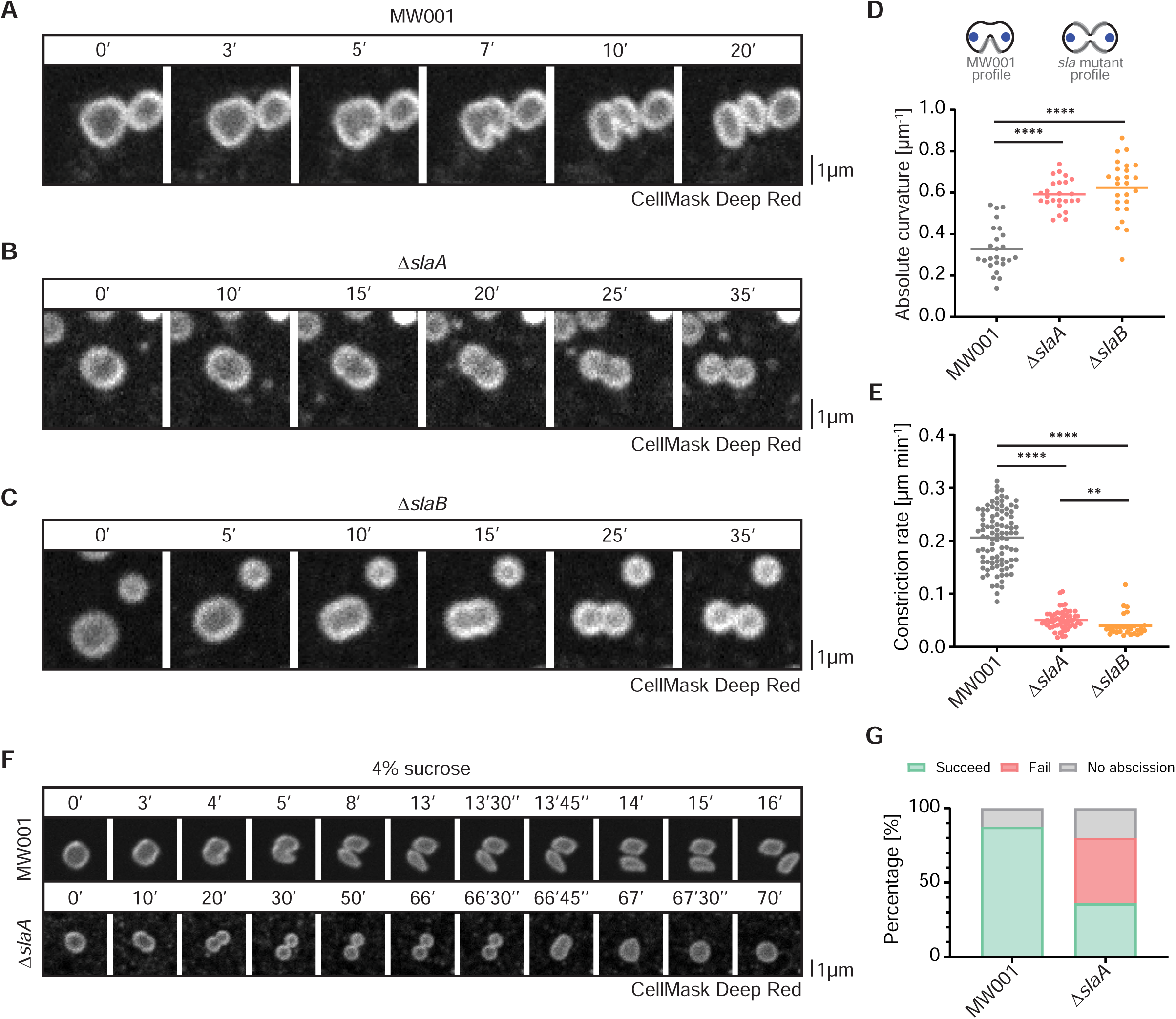
S-layer at the division bridge ensures proper constriction and abscission during cell division. Time-lapse imaging of (**A**) the uracil auxotrophic control strain MW001, (**B**) Δ*slaA* mutant and (**C**) Δ*slaB* mutant undergoing cell division. Deletion of either SlaA or SlaB results in changes in cell and division morphology, resulting in rounder cells. Representative division montages are shown. Cells were stained with CellMask Deep Red Plasma Membrane stain and imaged at 75°C in Brock medium supplemented with uracil. (**D**) Scatter plot with means indicated showing the quantification of the average curvature of the cell surface adjacent to the division bridge in MW001 and the S-layer mutants imaged under live conditions (MW001: 0.326 ±0.109μm^-1^; Δ*slaA*: 0.591 ±0.071μm^-1^; Δ*slaB*: 0.624 ±0.134μm^-1^). A diagram showing the profile used in this measurement is shown above. *P*-values were derived using Welch’s *t*-test (n=25; \*\*\*\**p* ≤0.0001; *n.s.*, no significance). (**E**) Scatter plot with means indicated showing the rate of constriction of MW001 and S-layer mutants (MW001: 0.2058 ±0.0527μm min^-1^; Δ*slaA*: 0.0507 ±0.0177μm min^-1^; Δ*slaB*: 0.0398 ±0.0210μm min^-1^). *P*-values were derived using Welch’s *t*-test (n ≥28; \*\*\*\**p* ≤0.0001; \*\**p* ≤0.01). (**F**) Time-lapse imaging of MW001 (top) and Δ*slaA* mutant (bottom) performed at 75°C in Brock medium supplemented with uracil, in the presence of 4% sucrose. Cells were stained with CellMask Deep Red Plasma Membrane stain. The Δ*slaA* mutant lacking an S-layer is more susceptible to stress, resulting in division failure. Shown here are representative montages. (**G**) Quantification of cell division failure in MW001 control cells and Δ*slaA* mutant dividing in 4% sucrose medium, imaged under live conditions (n ≥25; MW001: 87.5% succeed, 12.5% no abscission; Δ*slaA*: 36% succeed, 44% fail, 20% no abscission).

## Discussion

The self-assembly of large-scale protein structures has been studied in a host of organisms including the bacteria flagella (44), virus capsids (45), or even prions and amyloid fibrils in higher eukaryotes (46, 47). Such self-assembling 2D lattices also play a particularly important role in membrane remodelling in eukaryotic membrane trafficking, where the self-assembly of clathrin, COPI and COPII lattices coat membranes acts as a first step in vesicle formation (48–50). In these cases, the regulated formation of a passive, self-assembling protein coat (e.g. of Sec23/24, Sec13/31 (51), or clathrin (52)) generates positive membrane curvature. Energy-dependent processes (driven by the Sar1 GTPase, the Dynamin GTPase, or actin filament formation), then catalyse membrane scission (53, 54).

In a similar vein, in this paper we show how the energy-independent self-assembly of an SlaA lattice on membrane anchors on the surface of *Sulfolobus* cells assists in the remodelling of the membrane during cytokinesis. In the case of *Sulfolobus*, AAA-ATPase Vps4-dependent remodelling of composite ESCRT-III polymers then drives cytokinesis and abscission, leading to the formation of two daughter cells. In these examples, we can see how the collaboration of self-assembling polymers on the surface of a membrane, together with local force generation, lead to membrane deformation and scission. In the case of *Sulfolobus* division, the S-layer lattice is assembled as a coat on interphase cells that are near spherical, even though our data suggests that SlaA lattices prefer to bind to membranes that are flat. This suggests the possibility that there is stored elastic energy in the positively curved interphase lattice that is relieved during cytokinesis to help to form the furrow as a prelude to scission. This resembles what has been proposed to happen in eukaryotic cells when the mechanical energy stored in flat clathrin lattices on the plasma membrane assists in the formation of endosomes (55, 56). Despite this, the S-layer must be flexible, since in wildtype cells it completely cover the entire cell surface at all times – and completely coats small extracellular vesicles generated by *Sulfolobus* cells at division (34, unpublished data).

Importantly, there is no evidence of new S-layer insertion at the site of membrane furrowing in dividing Sulfolobus cells (Fig. S2D). Instead, the S-layer appears to form a stable lattice that grows as cells grow, and completely covers the division bridge, as shown using cryo-EM (Fig. 3G and S3G). Thus, the way the S-layer in *Sulfolobus* aids division appears to be a mechanical feature of the fully assembled lattice, not the result of dynamic local insertion – as is the case in Euryarchaea like Haloferax (14, 29), where the S-layer is laid down at the division plane to assist in division. In line with this, we did not observe any effects of the S-layer proteins on the localisation or dynamics of the ESCRT-III division ring (Fig. S4A and S4B). However, there was clear overlap in the patterns of CdvB1 recruitment to the internal side of the membrane in dividing Vps4^E209Q^ Δ*slaA* cells and the pattern of exogenously added SlaA (Fig. 3C-E). This suggests the possibility that SlaA lattices prefer membranes that are rigidified by an underlying ESCRT-III coat.

Taking together, these data lead us to propose that the S-layer flattens the membrane at the furrow on either side of the division bridge of dividing cells, where CdvB1 accumulates, to accelerate furrow ingression and scission (Fig. 4D and 4E). This fits with the observed preference of SlaA lattices for planar packing *in vitro* (20, 21). It also explains why in the absence of an S-layer, cells appear to divide by medial ring constriction, whereas control cells divide in a process that looks like slicing (Fig. 4A-C). The idea that the S-layer plays a simple mechanical role in cytokinesis is supported by its ability to protect cells subjected to mechanical shocks induced by changes in osmotic pressure from division failure (Fig. 4F and 4G). Although much harder to control, similar results were obtained when MW001 and *sla* mutant cells were subjected to mild mechanical compression.

In summary, our results show that S-layer monomers can self-assemble on the cell surface, in line with studies that have shown this unique ability *in vitro*, either in solution or on artificial substrates (57–61). This process of self-assembly generates a relatively rigid lattice that grows with the cell. Most strikingly of all, this structure both protects the bounding membrane of cells from environmental hazards, whilst also aiding their rapid and efficient division - two properties that one might have expected to be at odds with one another.

## Materials and Methods

### Strains, culture media and growth conditions

All *S. acidocaldarius* strains were grown at 75°C, pH 3.0, in Brock medium supplemented with 0.1% (w/v) NZ-amine and 0.2% (w/v) sucrose (62). Strains used in this study are found in Table 1. The uracil auxotroph strain MW001 and all deletion mutants were further supplemented with 4μg/ml uracil (35). Solid Brock medium was prepared by mixing pre-heated twice concentrated liquid Brock medium pH 5.0 supplemented with 0.2% NZ-amine (w/v), 0.4% sucrose (w/v), 20mM CaCl_2_, and 1.2% Gelrite (w/v), in a 1:1 ratio.

**Table 1:** Strains used in this study.

| Genotype | Strain | Figures |
| --- | --- | --- |
| *ΔpyrEF* | MW001 | 1A-C, 3A-B, 4A, 4D-G, S1C, S1F, S2A-D, S2F, S4A-B |
| *ΔpyrEF ΔslaA* | SF40 | 1A-C, 2F, 3A-B, 4B, 4D-G, S1A, S1F, S2F, S4A-B |
| *ΔpyrEF ΔslaB* | SF69 | 1A-C, 3A-B, 4C-E, S1A, S1F, S2F, S4A-B |
| *ΔpyrEF ΔslaAB* | SF101 | 1A-C, 3A-B, S1A, S1F, S2F, S4A-B |
| *ΔpyrEF Δsaci1846* | SF154 | 1C, S1A, S1E, S1F, S2F |
| *ΔpyrEF ΔslaB Δsaci1846* | SF146 | 1C, S1A, S1F, S2F |
| *ΔpyrEF pSVA-promoter<sup>ara</sup>:SlaA-HA* | SF57 | 2A, 2E-F, S1C |
| *ΔpyrEF ΔslaA pSVA-promoter<sup>ara</sup>:SlaA-HA* | SF58 | 2A, 2C-D, 2F |
| *ΔpyrEF pSVA-promoter<sup>ara</sup>:Vps4<sup>E209Q</sup>-His<sub>6</sub>* | SF93 | 3C, 3G, S3A-B, S3G |
| *ΔpyrEF pSVA-promoter<sup>ara</sup>:Vps4<sup>E209Q</sup>-His<sub>6</sub> ΔslaA* | SF85 | 3C-F, S3A-F |
| *ΔpyrEF pSVA-promoter<sup>ara</sup>:Vps4<sup>E209Q</sup>-His<sub>6</sub> ΔslaB* | SF86 | 3G, S3A-B, S3G |
| *ΔpyrEF pSVA-promoter<sup>ara</sup>:Vps4<sup>E209Q</sup>-His<sub>6</sub> ΔslaAB* | SF130 | S3A-B |
| *Wild-type* | DSM639 | S1B, S2E |
| *ΔpyrEF pSVA-promoter<sup>ara</sup>:HA-SlaB* | SF72 | S1C |

All liquid cultures were passaged up to 2 times and grown to exponential phase to an OD_595_ of 0.10-0.25 prior to use in all experiments, unless otherwise stated. Cells were fixed by three stepwise additions of 4°C ethanol to a final concentration of 70% and then stored at 4°C, as described previously (63). Induction of protein expression in overexpression strains and rescue mutants was performed by addition of L-arabinose to the culture medium to a final concentration of 0.2% (w/v) in exponentially growing cultures.

### Electrocompetent *S. acidocaldarius* cells

Electrocompetent cells were made by growing 50mL of the strain of interest to an OD_595_ of 0.20-0.30. The culture was cooled down on ice, followed by collection via centrifugation at 2500rcf for 10min at 4°C. The cell pellet was washed twice with 30mL of ice-cold 20mM sucrose. Next, the pellet was resuspended in 1mL of ice-cold 20mM sucrose and transferred to a fresh microcentrifuge tube. Cells were centrifuged at 2500rcf for 10min at 4°C, resuspended with ice-cold 20mM sucrose to a final theoretical OD_595_ of 10, split into 50μL aliquots and stored at -70°C.

### Molecular genetics

Deletion mutants were constructed as described previously, using the pSVA406 vector system (35). Briefly, the upstream and downstream regions of a gene of interest was cloned via restriction digest into pSVA406. Positive clones were identified via miniprep and analytical digestion. Plasmids of interest were methylated by transformation into *E. coli* ER1821, and subsequently purified using a GeneJET plasmid miniprep kit (ThermoFisher Scientific, K0503).

Transformation of electrocompetent *S. acidocaldarius* was performed following established protocols for *Sulfolobus* genetics (35). A 50μL aliquot of electrocompetent *S. acidocaldarius* was thawed on ice. 200ng of methylated plasmid DNA was next added and the mixture transferred to a 1mm Gene Pulser Electroporation Cuvette (BioRad, 1652089). Electroporation was performed at 2000V, 25μF and 600Ω. 400μL of pH 5.0 Brock medium without sucrose supplemented was then added and the cells incubated in a fresh microcentrifuge tube at 75°C for 60min for recovery. 150μL of cells were then plated on solid Gelrite-Brock plates for 5 days at 75°C. Transformed colonies were next selected and inoculated in 10mL of liquid Brock medium supplemented with 0.1% (w/v) NZ-amine and 0.2% (w/v) sucrose and incubated for 2 days at 75°C. Positive clones were identified through genomic DNA extraction and genotyping of the overlapping regions of the locus of interest. Genomic DNA extraction was performed by first pelleting 500μL of cells through centrifugation at maximum speed for 30s. Next, 20μL of 0.1M NaOH was added and the pellet dissolved, followed by addition of 80μL of 0.1M Tris-HCL pH 8.0.

20μL of strains containing the inserts were next streaked onto Gelrite-Brock plates supplemented with 4μg/ml uracil and 100μg/ml 5-fluoroorotic acid (5-FOA; Zymo Research, P9001-1). Following 5 days of incubation at 75°C, colonies were selected and incubated in 10mL of liquid Brock medium supplemented with 0.1% (w/v) NZ-amine, 0.2% (w/v) sucrose, 4μg/ml uracil and 100μg/ml 5-FOA for 2 days at 75°C. Positive clones were identified through genomic DNA extraction and genotyping of the overlapping regions of the locus of interest. Positive clones were frozen in Brock medium containing 50% glycerol (v/v) and stored at -70°C.

The SlaAB double deletion mutants was constructed by deletion of the entire operon from the start codon of SlaA (*saci2355*) to the stop codon of SlaB (*saci2354*) inclusive in electrocompetent MW001, using the protocol described above. The SlaB Saci1846 double deletion mutant was constructed by deletion of SlaB followed by a subsequent deletion of *saci1846* in electrocompetent Δ*slaB* cells.

Overexpression and rescue mutants were cloned into the pSVAaraHA-stop plasmid backbone, typically between the NcoI and XhoI restriction sites (64). Transformation and selection of electrocompetent *S. acidocaldarius* was performed as described above. The SlaA overexpression construct was created using two fragments, with a silent A-G point mutation introduced at DNA position 2079 via primer design to create an internal BamHI site for cloning between NcoI and BamHI, followed by BamHI and XhoI for the second fragment. This was done to avoid the endogenous NcoI site in the gene that is required for restriction cloning into the pSVAaraHA-stop plasmid. The HA-SlaB construct was procured commercially as a double stranded DNA fragment (gBlock, Integrated DNA Technologies) flanked by NcoI and XhoI restriction sites, with the HA tag (YPYDVPDYA) inserted after the signal sequence (amino acid residue S24) of SlaB. Vps4^E209Q^ dominant negative mutant overexpression construct was obtained from the lab collection (3).

### Whole genome sequencing

20mL of OD_595_ 0.25 cells were pelleted and genomic DNA extraction performed by phenol/chloroform extraction. Cell pellets were first resuspended in 250μL of 10mM Tris-HCl pH 8.0, 1mM EDTA, 150mM NaCl, 0.1% TritonX-100 (TENT) buffer and incubated at room temperature for 10min. Next, 250μL of phenol:chloroform:isoamyl alcohol (25:24:1, v/v) (Invitrogen, 15593031) was added and the mixture vortexed for 30s. The mixture was next centrifuged for 5min at 16,000rcf at room temperature. Following the transfer of the upper aqueous phase to a fresh microcentrifuge tube, the phenol/chloroform extraction was repeated twice. 200μL of the final upper aqueous phase was then transferred to a fresh microcentrifuge tube. Next, 150μL of 5M ammonium acetate and 875μL of 100% ethanol was added and this mixture was stored at -20°C overnight for DNA precipitation. The precipitated DNA was then pelleted at 16,000rcf for 5min, and the DNA pellet washed twice in ice-cold 70% ethanol. The ethanol was removed and the pellet air dried at room temperature for 10min. The DNA pellet was then resuspended in 50μL of nuclease free water and stored at 4°C.

DNA concentration was determined using a Qubit dsDNS BR assay (Invitrogen, Q33265) on a Qubit Flex Fluorometer (Invitrogen), following the manufacturer’s instructions. Whole genome sequencing and genome assembly was performed by Genewiz (Azenta Life Sciences).

### Cryo-EM and cryo-ET sample preparation, data collection and image analysis

Samples for cryo-EM and cryo-ET were prepared and performed as described previously (14, 65, 66). In summary, 2.5μL of the *S. acidocaldarius* cell culture were applied onto glow-discharged Quantifoil R3.5/1 Cu/Rh grids, blotted for 4-5 seconds with a blot force ranging from -10 to -4 after waiting 10 seconds and immediately plunge frozen into liquid ethane maintained at -178°C using a Vitrobot Mark IV (Thermo Fisher Scientific) cooled to 10°C with 100% chamber humidity.

Cryo-EM images were collected using a Glacios TEM (Thermo Fisher Scientific) operating at 200kV using a Falcon 3 (Thermo Fisher Scientific) direct electron detector. *S. acidocaldarius* MW001, Δ*slaA*, Δ*slaB* and Δ*slaAB* were imaged at a nominal magnification of 73,000 resulting in a 1.994Å pixel size, using a total dose of 40e^-^/Å^2^ within a defocus range of -6 to -10μm. Δ*slaAB,* Δ*saci1846* and Δ*saci1846* Δ*slaB* were imaged at a nominal magnification of 92,000 resulting in a 1.583Å pixel size, using a dose of 40e^-^/Å^2^ within a defocus range of -6 to -10μm. Vps4^E209Q^ Δ*slaA* with exogenously added SlaA were imaged at a nominal magnification of 57,000 resulting in a 2.545Å pixel size using a dose of 40e^-^/Å^2^ within a defocus range of -6 to -10μm. All electron cryomicrographs shown have been lowpass filtered to improve visibility of the cell envelope and S-layer.

Tilt-series data was acquired using a Krios G3 TEM (Thermo Fisher Scientific) operating at 300kV using a K3 (Gatan) direct electron detector equipped with a Quantum energy filter (Gatan) with a slit width of 20eV. Data collection was performed using SerialEM software (67) with the dose-symmetric scheme (68) at a nominal magnification of 26,000, resulting in a pixel size of 2.809Å in counting mode. Tilt images were acquired at a tilt range of ±60° with 3° tilt increment with a total dose of 121-122e^-^/Å^2^ and with defoci ranging from -4 to -7μm (Vps4^E209Q^) or -7 to -11μm (Vps4^E209Q^Δ*slaB*).

Frames from tilt series movies were motion corrected using RELION (69) and tilt-images were aligned and SART filtered using AreTomo (70). Final tomograms were reconstructed using the RELION5 pipeline (71) with 4x binning resulting in a final pixel size of 11.236Å.

### Immunofluorescence labelling

Fixed cells were first pelleted via centrifugation at 8000rcf for 3min to remove ethanol and washed twice in 1mL of phosphate-buffered saline supplemented with 0.2% Tween-20 (Sigma-Aldrich, P9416) and 3% Bovine Serum Albumin (Sigma-Aldrich, A7030) (PBSTA) for rehydration and blocking. Cells were then incubated overnight at room temperature in 200μL of PBSTA containing primary antibodies (Table 2) with 400rpm agitation on an Eppendorf thermomixer. Following two washes in 1mL phosphate-buffered saline supplemented with 0.2% Tween-20 (PBST), 200μL of PBSTA containing secondary antibodies (Table 2) were added for 2h at room temperature and 400rpm agitation on an Eppendorf thermomixer. Labelling of cell surface was performed by incubating with 5μg/ml concanavalin A conjugated to Alexa Fluor 647 (Thermo Fisher Scientific, C21421) during secondary antibody incubation. Following two washes in 1mL PBST, cells were resuspended in 200μL of PBST containing either 1μg/mL DAPI (4′,6-diamidino-2-phenylindole; Thermo Fisher Scientific, 62248) for imaging, or 2μM Hoechst (Thermo Fisher Scientific, 62249) for flow cytometry experiments.

**Table 2:** Antibodies used in this study.

| Antibody | Host organism | Dilution | Catalog number |
| --- | --- | --- | --- |
| Anti-HA IgG | Mouse | 1:1000 | Invitrogen, 261831MG |
| Anti-CdvB serum | Mouse | 1:1000 | - |
| Anti-CdvB1 IgY | Chicken | 1:1000 | - |
| Anti-CdvB2 peptide Ab | Guinea pig | 1:1000 | - |
| Anti-Mouse IgG, AF488 | Goat | 1:1000 | Invitrogen, A11034 |
| Anti-Chicken IgY, AF546 | Goat | 1:1000 | Invitrogen, A11040 |
| Anti-Guinea pig IgG, AF647 | Goat | 1:1000 | Invitrogen, A21450 |
| Anti-Mouse IgG IRDye 800CW | Goat | 1:10,000 | LI-COR Bio, 926-32210 |

### Microscopy

Fixed cells were imaged in Lab-Tek chamber slides (Thermo Fisher Scientific, 177437PK) coated with 2% polyethyleneimine for 2h at 37°C. Chambers were rinsed with Milli-Q water twice before addition of 200μL of stained cells. Cells were immobilized by centrifugation for 1h at 750rcf by taping the chamber slides down to a centrifuge microplate adaptor. Imaging was performed using a Nikon Eclipse Ti2 inverted microscope equipped with a Yokogawa SoRa scanner unit and Prime 95B scientific complementary metal-oxide semiconductor (sCMOS) camera (Photometrics). All images were acquired using a 100x oil immersion objective (Apo TIRF 100×/1.49, Nikon) with immersion oil (Nikon, Type F2). A further 2.8x magnification was achieved using the 2.8x lens in the SoRa unit. z-steps were set at 0.22μm. All analysis and z-axis projections were performed in ImageJ. Quantification of ConA intensity was performed in ImageJ using the segmented line function with a line width of 2 pixels.

### Flow cytometry

Fixed cells were stained as described above and DNA labelled with 2μM Hoechst. Flow cytometry experiments were performed on a BD Biosciences LSRFortessa. Laser excitation wavelengths of 355, 488, 561 and 633nm were used in conjunction with the emission filters 450/50, 530/30, 586/15 and 670/14 respectively. Both side and forward scatter information was recorded as well. All analysis of flow cytometry data was performed on FlowJo v10 software.

### Labelling of cell surface with NHS ester dyes

30mL of exponential growing *S. acidocaldarius* cells were first collected via centrifugation at 4000rcf for 5min. The cell pellet was washed once in 1mL of 50mM HEPES pH 8.0 buffer before being resuspended in 1mL of 50mM HEPES pH 8.0. Pulse labelling was performed by addition of 3μL of a 1mg/mL stock of Alexa Fluor 488 NHS succinimidyl ester dye (Invitrogen, A20000) in DMSO to the solution and incubated in an Eppendorf thermomixer at room temperature for 30min with 500rpm shaking. Next, the cells were washed once in 1mL of 50mM HEPES pH 8.0 and once in 1mL of pre-warmed Brock medium. The cell pellet was resuspended in Brock medium and diluted into 25mL of pre-warmed Brock medium and allowed to grow for 2h. The process was then repeated with Alexa Fluor 555 NHS succinimidyl ester dye (Invitrogen, A20009) for chase labelling.

Quantification of pulse and chase labelling, as well as labelling optimization, was performed in ImageJ using the segmented line function with a line width of 2 pixels. Line plots were generated in ImageJ and the raw values exported for processing in Microsoft excel and Graphpad Prism 7 software.

Optimization of labelling with NHS ester dyes was performed as described above, using Alexa Fluor 488 NHS succinimidyl ester dye (Invitrogen, A20000) in different buffers. 50mM Citrate buffer at pH 5.0 or pH 6.0, or 50mM HEPES buffer at pH 7.0 or 8.0 was used for the wash steps and incubation with the dye. Further analysis of NHS ester dye labelled proteins was performed via SDS-PAGE. 10mL of OD595 0.1 of labelled cells was collected via centrifugation and resuspended in 200μL of 1x NuPAGE LDS sample buffer (Thermo Fisher Scientific, NP0007) containing 10% β-mercaptoethanol and incubated at 98°C for 5min. Following a brief centrifugation, samples were loaded and run on a NuPAGE 4 to 12% Bis-Tris gels (Invitrogen, NP0321BOX) at a constant 150V for 60-80min using MOPS running buffer. The gel was imaged directly on a BioRad ChemiDoc MP imaging system using the Alexa 488 function (UV 520/30).

### Proteinase K treatment

30mL of exponentially growing SlaA-HA and HA-SlaB overexpression strains were induced with arabinose for 4h. Cells were then pelleted via centrifugation at 4000rcf for 5min. Cell pellets were transferred to a clean microcentrifuge tube and washed twice in 50mM HEPES buffer pH 8.0. The pellet was then resuspended in 1mL of 50mM HEPES buffer pH 8.0, and 20μL of 20mg/mL proteinase K added. This was incubated in an Eppendorf thermomixer at 60°C for 30min with 500rpm shaking. Next, proteinase K was removed by washing the cells once in 1mL of 50mM HEPES buffer pH 8.0 and once in 1mL of pre-warmed Brock medium. This was then diluted into 25mL of pre-warmed Brock medium and allowed to grow for 2 to 4h. The cells were next fixed by three stepwise addition of 4°C ethanol to a final concentration of 70% and then stored at 4°C, as described previously.

### S-layer purification

1L of wild-type *S. acidocaldarius* DSM639 was grown to an OD_595_ of 0.8, following standard cell husbandry protocols as described above. Cells were centrifuged at 4000rcf for 30min using four 250mL large volume centrifuge tubes (Corning, CLS430776) and the supernatant was discarded. The cell pellets were resuspended in water and transferred to two 50mL falcon tubes before being centrifuged at 4000rcf for 30min. The supernatant was discarded and the falcon tubes containing the cell pellets were stored at -70°C until further use.

S-layer purification was performed as described previously (21, 38), with minor modifications for compatibility with benchtop centrifugation. Each cell pellet was first thawed and resuspended in 40mL of 10mM HEPES buffer pH 7.0 containing 2mM EDTA and 1mM PMSF (HEPES wash buffer), supplemented with 600μL of 10% SDS to obtain a final concentration of 0.15% SDS. 100μg/μL DNaseI (ITW reagents, A3778.0100) and 4mM MgCl_2_ was added and the mixture incubated for 1h at 37°C. A further 8mL of 10% SDS was added to bring the final concentration to 2% and the mixture left overnight on a roller mixer at room temperature.

The mixture was next centrifuged for 1h at room temperature at 4000rcf and the supernatant discarded. The pellet was washed with 30mL of HEPES wash buffer +2% SDS for 1h at 75°C with 200rpm shaking in the incubator, followed by centrifugation for 1h at room temperature at 4000rcf and the supernatant discarded. This wash was repeated for a further 2 times, before a final resuspension in 30mL of HEPES wash buffer +2% SDS overnight. The following day, the mixture is briefly heated at 75°C for 15min to dissolve any precipitated SDS, before centrifugation for 1h at room temperature at 4000rcf. The pellet was then washed thrice in 30mL milli-Q water, before transferring to a clean microcentrifuge tube. A further 3 washes in 1mL of milli-Q water with centrifugation for 15min at 20,000rcf is performed, before the pellet is solubilised in 1mL of 20 mM Na-carbonate buffer pH10 at 75°C for 30min.

Successful purification of S-layer proteins was checked by running on an SDS-PAGE gel and staining with InstantBlue Coomassie Protein Stain (Abcam, ab119211). Concentration of purified proteins was estimated via spectrophotometric quantitation at 280nm and calculated using Beer–Lambert law, assuming SlaA with a theoretical molar extinction coefficient (72) of 1.175 as the major component of the solution. Further validation of purified proteins was performed via mass spectrometry analysis of the solution, which confirmed the presence of SlaA and SlaB.

### S-layer in vivo assembly

30μg of purified S-layer proteins were diluted 1:1(v/v) with 50mM HEPES pH 7.0. Next, 1μL of a 1mg/mL stock of Alexa Fluor 488 NHS succinimidyl ester dye (Invitrogen, A20000) in DMSO was added and the mixture incubated for 15min at room temperature with gentle agitation. 3mL of cells in exponential growth phase were then transferred to a 15mL falcon tube, and labelled S-layer proteins added. This was incubated at 75°C with shaking for 1h. The cells were next fixed by three stepwise addition of 4°C ethanol to a final concentration of 70% and then stored at 4°C, as described previously. For cryo-EM analysis of S-layer in vivo assembly, unlabelled S-layer proteins were added instead.

### ConA pull-down and Western blotting

25mL of OD_595_ 0.2 cultures of MW001, SlaA-HA and HA-SlaB cells were collected via centrifugation at 10,000rcf. Cell pellets were resuspended in 300μL of IP lysis buffer (25mM HEPES pH 7.5, 10mM KCL, 1mM CaCl_2_, 1mM MnCl2) supplemented with 1x RIPA Lysis buffer (Merck, 20-188) and 1x cOmplete protease inhibitor (Roche, 11836170001). The cells were next lysed in a Bioruptor Plus sonication device at 4°C for 7 cycles of 30s on and 30s off, at low power setting. Lysed cells were then centrifuged for 5min at 10,000rcf at 4°C to pellet cellular debris. 200μL of the supernatant were transferred to fresh microcentrifuge tubes and diluted with 300μL of IP lysis buffer. 80μL of the remaining supernatant of each sample was transferred to a fresh tube and 20μL of 1x NuPAGE LDS sample buffer (Thermo Fisher Scientific, NP0007) containing 10% β-mercaptoethanol added, and set aside for use as cell lysate control.

60μL of magnetic agarose Concanavalin A Beads (Antibodies-online, ABIN6952467) were equilibrated by washing thrice in 1mL of IP lysis buffer on a magnetic rack. Pre-equilibrated beads were resuspended in IP lysis buffer and equal amounts added to each prepared supernatant earlier. ConA pull-down was performed on a rotating mixer at 4°C overnight. Following that, The supernatant was discarded and beads washed thrice with 1mL of ice-cold IP lysis buffer for 5min on the rotating mixer at 4°C. Next, the beads were resuspended in 60μL of 1x NuPAGE LDS sample buffer (Thermo Fisher Scientific, NP0007) containing 10% β-mercaptoethanol. Cell lysate control and ConA pull-down samples were incubated at 98°C for 5min. Following a brief centrifugation, the samples were loaded and ran on a NuPAGE 4 to 12% Bis-Tris gels (Invitrogen) at a constant 150V for 60-80min using MOPS running buffer. Proteins were transferred to nitrocellulose membranes at a constant 100V at 4°C for 60min. Blocking was performed using 5% milk in PBST for 1h, followed by incubation with primary antibodies (Table 2) in PBST +5% milk overnight at 4°C with gentle agitation. The membrane was washed in PBST thrice for 5min and then incubated with PBST +5% milk containing IRDye secondary antibodies (Table 2) for 2h. Following three washes in PBST, the membrane was imaged using a Bio-Rad ChemiDoc system.

### Mass spectrometry

ConA IP beads were resuspended in 50μL of 25mM AMBIC 0.1% RapiGest and digested for protein identification by mass spectrometry. Briefly, cysteines were reduced by addition of 5μL of a 11.1mg/mL DTT solution to achieve a final concentration of 4mM, and mixed via vortexing before incubating for 10min at 60°C. Alkylation was next performed by addition of 5μL of iodoacetamide (46.6mg/mL) to achieve a final concentration of 14mM, and incubated at room temperature in the dark for 30min. 1μg of trypsin was added for in-bead digestion for 3h at 37°C at 450rpm shaking on a Eppendorf thermomixer. Beads were removed and digestion performed overnight at 37°C. The following day, 1μL of trifluoracetic acid was added to a final concentration of 0.5% (v/v) and incubated for 45min at 37°C to stop digestion and induce RapiGest degradation. The mixture was centrifuged for 15min at 13,000rcf at 4°C. The supernatant was transferred to a fresh microcentrifuge tube and 5μL injected over a 60min programme for mass spectrometry analysis.

LC-MS/MS was performed on an Ultimate U3000 HPLC (ThermoFisher Scientific) hyphenated to an Orbitrap QExactive Classic mass spectrometer (ThermoFisher Scientific). Peptides were trapped on a C18 Acclaim PepMap 100 (5µm, 300µm x 5mm) trap column (ThermoFisher Scientific) and eluted onto an IonOpticks Aurora Ultimate column (1.7µm, 75µm x 250mm) using 52 minute gradient of acetonitrile (7 to 40%). For data dependent acquisition, MS1 scans were acquired at a resolution of 70,000 (AGC target of 1e6 ions with a maximum injection time of 65ms) followed by ten MS2 scans acquired at a resolution of 17,500 (AGC target of 2e5 ions with a maximum injection time of 100ms) using a collision induced dissociation energy of 25. Dynamic exclusion of fragmented m/z values was set to 40s.

Raw data were imported and processed in Proteome Discoverer v3.1 (Thermo Fisher Scientific). The raw files were submitted to a iterative database search using Proteome Discoverer with Sequest HT and Inferis Rescoring against the UniProt reference proteome for *S. acidocaldarius* with methionine oxidation and cysteine carbamidomethylation set as variable and fixed modifications, respectively. The spectra identification was performed with the following parameters: MS accuracy of 10ppm MS/MS accuracy of 0.02Da; up to two trypsin missed cleavage sites were allowed. Only rank 1 peptide identifications of high confidence (FDR<1%) were accepted.

### Trichloroacetic acid precipitation of SlaA from growth medium

*S. acidocaldarius* cells were grown to an OD_595_ of 0.5 under standard culturing conditions. 20mL of each culture was centrifuged at 4000rcf for 10min to pellet the cells. 15mL of the supernatant is then removed carefully, and passed through a 0.22μm syringe filter into a fresh tube to remove all remaining cells.

555μL of 100% (w/v) trichloroacetic acid (Sigma-Aldrich, T6399) was added to 5mL of the filtered media to a final concentration of 10% (v/v) and the mixture incubated on ice for 1h with intermittent mixing. Precipitated proteins were pelleted via centrifugation at 20,000rcf at 4°C for 10min. The pellet is next washed in 1mL of ice-cold acetone, and pelleted again at 20,000rcf at 4°C for 10min. The acetone was discarded and the pellet dried in a Eppendorf Vacufuge plus vacuum concentrator for 5min at room temperature. Precipitated proteins were then resuspended in 50μL of 1x NuPAGE LDS sample buffer (Thermo Fisher Scientific, NP0007) containing 10% β-mercaptoethanol, and incubated at 98°C for 5min. 20μL of each sample was loaded and run on a NuPAGE 4 to 12% Bis-Tris gels (Invitrogen, NP0321BOX) at a constant 150V for 60-80min using MOPS running buffer. Proteins were visualised via staining of the gel with InstantBlue Coomassie Protein Stain (Abcam, ab119211).

### Live cell microscopy

Live cell imaging was performed using the Sulfoscope set-up as described previously (3, 34, 63). Briefly, Attofluor chambers (Invitrogen, A7816) were assembled with 25mm coverslips and filled with 300µL Brock media supplemented with 0.1% (w/v) NZ-amine and 0.2% (w/v) sucrose. The media was allowed to dry onto the surface of the coverslip at 75°C before the chambers were rinsed twice with fresh Brock media and placed into the pre-heated Sulfoscope. The setup was allowed to equilibrate to 75°C. 5mL of *S. acidocaldarius* cell culture at an OD_595_ of 0.15-0.30 was stained with 1:5000 CellMask Deep Red Plasma Membrane (Invitrogen, C10046) and kept at 75°C in a polystyrene box with pre-warmed metallic beads (Gibco Lab Armor, 10120988).

Gelrite-Brock medium pads were prepared by cutting from a petri dish with a 7mm diameter circle punch and placed onto 13mm circular coverslips. Pads were prewarmed for 5min at 75°C for 5min in a bead bath prior to imaging. 400µL of the stained cell suspension was then transferred to the pre-heated Attofluor chamber and immobilised using Gelrite-Brock medium pads with the concave edges of the pad in the middle of the imaging chamber. Imaging was performed at this concave edge where diffusion is limited but cells are not subjected to mechanical stress by the pad.

Images were acquired as described above, with a 60× oil immersion objective (Plan Apo 60×/1.45, Nikon) using a custom formulated immersion oil for high temperature imaging (maximum refractive index matching at 70°C, n=1.515 ±0.0005; Cargille Laboratories). Images were acquired at intervals of 15s for 2.5h or until any cell death was observed. XY drift was corrected after acquisition using the ImageJ plugin StackReg (73).

### Curvature measurements

Curvature measurements were performed using the Kappa plugin on Fiji (74). Data points were exported and analysed on Microsoft Excel and GraphPad Prism10. Spearman’s rank correlation coefficient was calculated between the absolute curvature and exogenous S-layer labelling fluorescence intensity to determine the relationship between the two variables. Quantification in live cells was obtained by averaging the curvatures on the regions adjacent to the division bridge in late constriction cells.

### Statistical analysis

All statistical analysis was performed on Microsoft Excel or GraphPad Prism 10 software. All data presented are shown as mean ±SD, unless otherwise specified. Significance was defined as *p* ≤0.05. Significance levels used were \**p* ≤0.05, \*\**p* ≤0.01, \*\*\**p* ≤0.001 and \*\*\*\**p* ≤0.0001.

## Author contributions

All cell biology, genetics and biochemistry experiments were performed by SF. Live cell imaging, constriction rates and division failure analyses were performed by AC. Cryo-EM experiments and analyses were performed by IC and TAMB. BB supervised the project and co-wrote the manuscript with SF.

## Acknowledgements

Mass spectrometry analysis was/were performed at the Biological Mass Spectrometry and Proteomics Facility of the Medical Research Council Laboratory of Molecular Biology, Cambridge, UK. The authors would like to thank Dr Catarina Franco and Dr Farida Begum for the help with data analysis and helpful discussions on experimental design. The authors acknowledge the LMB electron microscopy facility for supporting sample preparation and data collection. The authors thank Jake Grimmett, Toby Darling and Ivan Clayson of the LMB Scientific Computing.

SF was supported by the Wellcome Trust (222460/Z/21/Z); IC was supported by European Molecular Biology Organization (EMBO) Long-Term Fellowships (ALTF 92-2022); AC was supported by a Marie Skłodowska-Curie individual fellowship (101068523) provided by UK Research and Innovation. This work was supported by the Human Frontiers Science Program (RGY0074/2021) awarded to TAMB, the Medical Research Council, as part of United Kingdom Research and Innovation (MC_UP_1201/31). TAMB would like to thank the European Molecular Biology Organization, the Wellcome Trust (225317/Z/22/Z) and the Leverhulme Trust. BB received support for work in *Sulfolobus* from the Medical Research Council-Laboratory of Molecular Biology (MC_UP_1201/27), the Wellcome Trust (222460/Z/21/Z). This work was supported by the Moore-Simons Project on the Origin of the Eukaryotic Cell, Simons Foundation (735929LPI).

## Data and materials availability

All data are available in the manuscript or the supplementary materials. All materials are available from BB or SF on request.

## Supplementary Figure Legends

**Figure S1.**
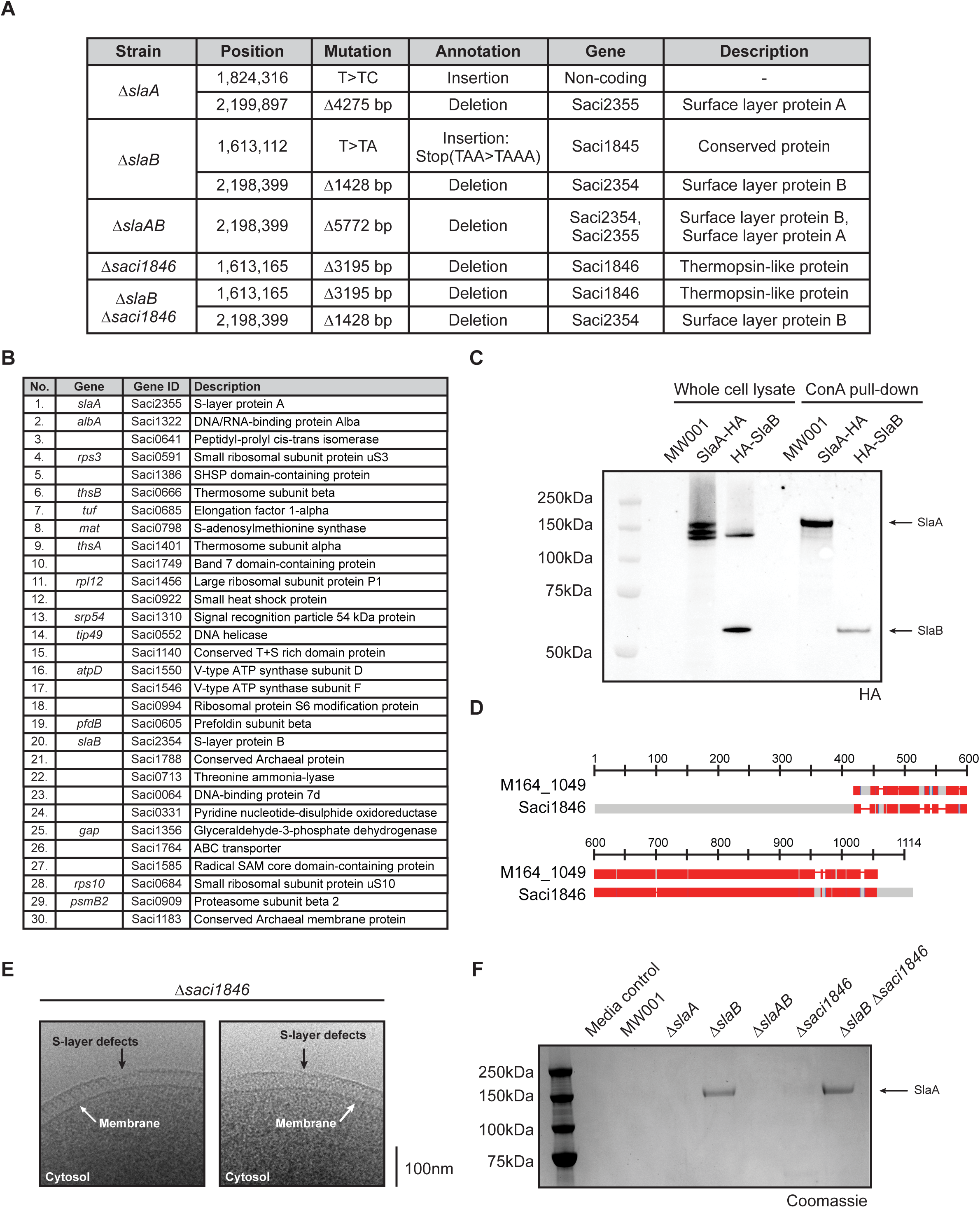
Validating S-layer mutants and their visualization with Concanavalin A. (**A**) Whole genome sequencing of the uracil auxotrophic background strain MW001 and the respective mutants generated in this study was performed. A summary of mutations identified compared to MW001 is presented. (**B**) Top 30 hits from mass spectrometry analysis of proteins pulled down from wild-type *S. acidocaldarius* DSM639 lysates using ConA magnetic beads. SlaA is the most abundant protein detected, and SlaB is also present in the pull-down. (**C**) Whole cell lysates of MW001 and HA-tagged SlaA or SlaB cells were incubated with magnetic ConA beads to assess ConA binding to the highly glycosylated S-layer proteins. Western blot of input cell lysates and the ConA pulldown assay shows successful pull-down of the HA-tagged SlaA and SlaB proteins, visualised using anti-HA antibodies. (**D**) Protein sequence alignment of M164_1049 and Saci1846 using the NCBI Multiple Sequence Alignment Viewer. Alignment positions are coloured red indicating highly conserved positions, blue indicating lower conservation and grey indicating unaligned residues. (**E**) Minor S-layer defects are observed in the Δ*saci1846* mutants, as shown in these representative cryo-EM images. (**F**) Coomassie-stained gel of TCA precipitated proteins from the growth medium after culturing of the indicated strains. SlaA can be detected in the medium of Δ*slaB* and Δ*slaB* Δ*saci1846* double mutants, showing the stripping of the SlaA layer from the cell surface.

**Figure S2.**
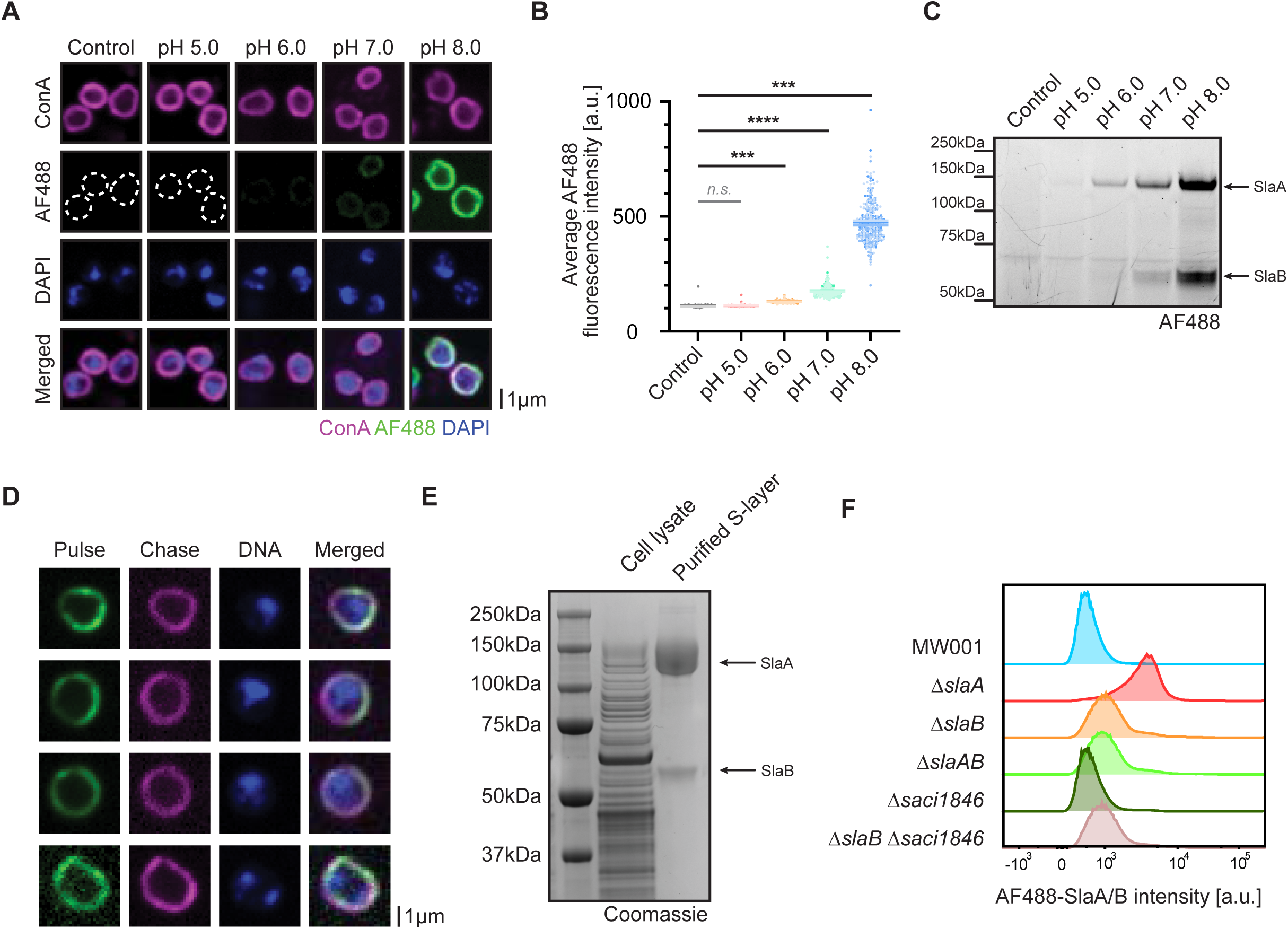
Labelling of the *S. acidocaldarius* cell surface with NHS ester dyes and exogenous S-layer proteins. (**A**) MW001 cells were labelled with Alexa Fluor 488 (AF488) NHS ester in Brock medium (control, pH 3.0), or at a range of different pH, showing strong labelling under basic conditions. Shown here are representative immunofluorescence images. (**B**) Quantification of labelling efficiency at different pH was performed by measuring average fluorescence intensities at the cell surface (N=3, n=100 cells each). Each biological replicate is represented by a different shade of colour and the means indicated (Control: 110.3 ±1.51; pH 5.0: 111.3 ±0.41; pH 6.0: 129.7 ±1.41; pH 7.0: 173.9 ±2.15; pH 8.0: 472.7 ±8.42). *P*-values were derived using Welch’s *t*-test (N=3, \*\*\**p* ≤0.001; \*\*\*\**p* ≤0.0001; *n.s.*, no significance). (**C**) SDS-PAGE gel of whole cell fractions following labelling at different pH, showing strong labelling at basic pH of two major protein bands corresponding to the molecular weights of SlaA and SlaB. Labelled proteins were visualised using the Alexa 488 function (UV 520/30) on a BioRad ChemiDoc MP imaging system. (**D**) MW001 cells were pulse labelled using AF488 (green) and chase labelled using Alexa Fluor 555 (magenta) NHS esters based dyes. The pulse-chase experiment indicates that S-layer biogenesis is not localised to a defined location on the cell surface. Shown here are representative cells including a dividing cell (bottom row). (**E**) Coomassie-stained gel showing purified wild-type SlaA and SlaB extracted from wild-type *S. acidocaldarius* DSM639 cultures. (**F**) Purified S-layer proteins labelled with AF488 NHS ester dye were incubated with S-layer mutants. Shown here are representative flow cytometry histograms of AF488-SlaA/B intensities in each mutant background, representing the efficiency of labelling these mutants with exogenous S-layer proteins. Only the Δ*slaA* mutant lacking the S-layer but containing the membrane anchors SlaB and Saci1846 is efficiently labelled with exogenous S-layer proteins (N=3, n=2.5×10^5^ events).

**Figure S3.**
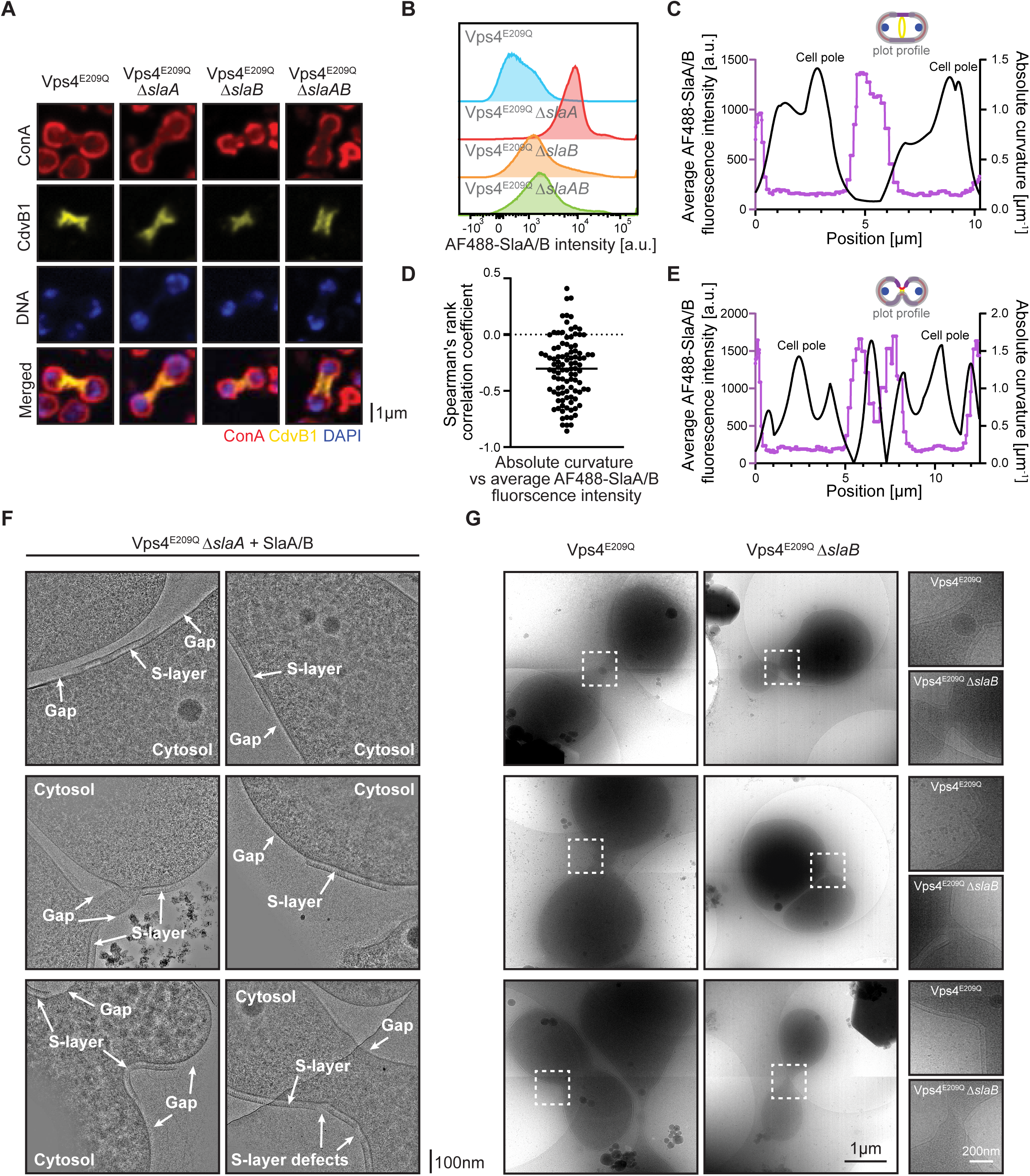
Labelling of S-layer mutants expressing Vps4^E209Q^ arrested at cell division with exogenous S-layer proteins. (**A**) Representative single-plane spinning-disk confocal images of MW001 and the S-layer mutants expressing the Walker B dominant negative mutant Vps4^E209Q^ shows cells arrested at division. (**B**) Purified S-layer proteins labelled with Alexa Fluor 488 (AF488) NHS ester dye were incubated with MW001 or the S-layer mutants overexpressing Vps4^E209Q^. Shown here are representative flow cytometry histograms of AF488-SlaA/B intensities in each mutant background, representing the efficiency of labelling these mutants with exogenous S-layer proteins. Only the Vps4^E209Q^ Δ*slaA* mutant lacking the S-layer but containing membrane anchors is efficiently labelled with exogenous S-layer proteins (N=3, n=2.5×10^5^ events). (**C**) Representative Kappa curvature analysis of a pre-constriction Vps4^E209Q^ Δ*slaA* cell showing the absolute point curvature (black) and the fluorescence intensity of exogenous S-layer protein (magenta) on the cell surface. A diagram showing the plot profile used indicated in grey is shown above. (**D**) Spearman’s correlation coefficient between the absolute point curvature and the fluorescence intensity of exogenous S-layer proteins on the surface of Vps4^E209Q^ Δ*slaA* cells (n=100 cells). The negative correlation suggests that the presence of exogenous S-layer proteins is negatively correlated with curved surfaces. (**E**) Representative Kappa curvature analysis of a late constriction Vps4^E209Q^ Δ*slaA* cell showing the absolute point curvature (black) and the fluorescence intensity of exogenous S-layer protein (magenta) on the cell surface. A diagram showing the plot profile used indicated in grey is shown above. (**F**) Gallery of representative cryo-EM images of Vps4^E209Q^ Δ*slaA* mutants with exogenous unlabelled S-layer proteins added. Exogenous S-layer proteins form lattices on the cell surface via self-assembly. Defects and gaps were observed at regions of high local curvature changes. (**G**) Gallery of representative cryo-EM images of Vps4^E209Q^ and Vps4^E209Q^ Δ*slaB* mutants. The partial S-layer of Δ*slaB* mutants are observed to preferentially localize to the division bridges. Magnified images of the division bridges indicated by the white boxes in each example are shown on the right.

**Figure S4.**
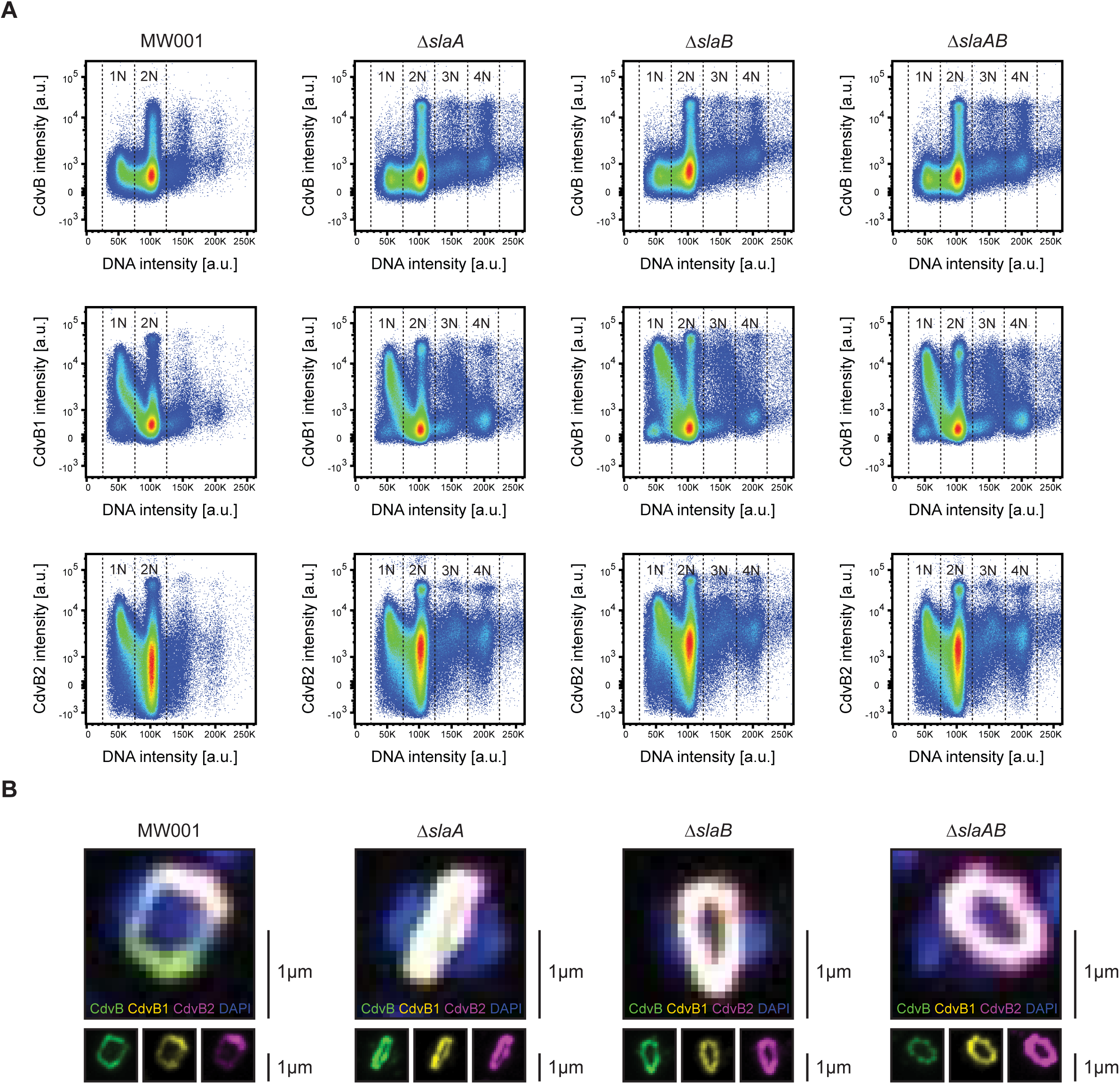
The ESCRT-III rings of S-layer mutants. (**A**) Representative flow cytometry scatter plots of asynchronous cultures of MW001 and the S-layer mutants. CdvB (top), CdvB1 (middle) and CdvB2 (bottom) signals are plotted against DNA in each strain. Each spot represents a single cell, with the density gradient going from blue to red. N=3, n=5.0×10^5^ events. (**B**) Representative maximum intensity z-axis projections of MW001 and the S-layer mutants during mitosis shows formation of the ESCRT-III rings. Cells were stained for DNA with DAPI (blue), CdvB (green), CdvB1 (yellow) and CdvB2 (magenta). No obvious changes in the morphology of the ring structures was observed in the S-layer mutants.

## Supplementary Movies Legends

**Supplementary Movie SM1. Cryo-ET of MW001 expressing Vps4^E209Q^.** The S-layer is present with no gaps on the cell surface of MW001 expressing the Walker B dominant negative mutant Vps4^E209Q^ arrested at division. Scale bar represents 100nm.

**Supplementary Movie SM2. Cryo-ET of MW001 expressing Vps4^E209Q^.** The S-layer is present with no gaps on the cell surface of MW001 expressing the Walker B dominant negative mutant Vps4^E209Q^ arrested at division. Scale bar represents 100nm.

**Supplementary Movie SM3. Cryo-ET of Δ*slaB* mutants expressing Vps4^E209Q^.** The S-layer patches present in Δ*slaB* mutants preferentially localizes to the division bridge in Δ*slaB* cells expressing the Walker B dominant negative mutant Vps4^E209Q^ arrested at division. Scale bar represents 100nm.

**Supplementary Movie SM4. Cryo-ET of Δ*slaB* mutants expressing Vps4^E209Q^.** The S-layer patches present in Δ*slaB* mutants preferentially localizes to the division bridge in Δ*slaB* cells expressing the Walker B dominant negative mutant Vps4^E209Q^ arrested at division. Scale bar represents 100nm.

